# Architecture of a pentameric assembly of the tail tube protein in SPR phages

**DOI:** 10.1101/2024.11.10.622895

**Authors:** Lin Wang, Yuhang He, Kaixiang Zhu, Sheng Cui, Xiaopan Gao, Kun Shang, Hongtao Zhu

**Affiliations:** Beijing National Laboratory for Condensed Matter Physics and Laboratory of Soft Matter Physics, Institute of Physics, Chinese Academy of Sciences, 100190, China; University of Chinese Academy of Sciences, Beijing, 100049, China; Songshan Lake Materials Laboratory, Dongguan, Guangdong, China; Medical School, Yan’an University, Yan’an, Shaanxi, China; NHC Key Laboratory of Systems Biology of Pathogens, National Institute of Pathogen Biology, Chinese Academy of Medical Sciences and Peking Union Medical College, Beijing, China

## Abstract

The majority of phages, viruses that infect prokaryotes, deliver their genomic material into host cells via a tail structure. Previous research has established that the central tube of these phages, composed of tail tube protein (TTP), is structurally conserved, typically forming either hexameric or trimeric rings. In this study, we revealed a novel pentameric assembly of TTP and present two cryo-EM structures at resolutions of 3.5 Å and 3.7 Å, respectively. Our detailed structural analysis demonstrates that the inner surface of the pentameric tube is highly negatively charged. Critical residues located on the loop between β3 and β4 play pivotal roles in the formation of the pentameric rings, and mismatches of the interactions between the stacked layers can induce the curvature of the tube. The cryo-EM structure of the TTP polymer at the tube’s end uncovered that the β-strands that span amino acids 27-65 move towards the central tunnel, potentially blocking the tunnel through which the phage genome is released. Our study provides new structural insights into a novel assembly of TTP, which will extend the understanding of phage assembly.

## INTRODUCTION

Phages are capable of infecting prokaryotes.^[1-3]^ The structure of a typical phage comprises several essential components for infection: an icosahedral protein shell, or head, that houses the phage’s genetic material, usually double-stranded DNA, and a tail structure that includes a tail sheath, tail tube, and base plate.^[4, 5]^ The tail can vary in length and may be flexible or rigid, contractile or noncontractile.^[5]^ Phages are classified into three families based on tail morphology: Siphoviridae (long flexible tail), Myoviridae (long contractile tail), and Podoviridae (short tail).^[4-7]^ The phage tail is a compelling subject of study due to its assembly pathway, structural characteristics, host recognition mechanisms, and role in cell wall perforation.^[7]^ The tail tube is typically formed by stacking hexameric rings of the tail tube protein (TTP) which consists of a sandwich of two antiparallel β-sheets, an α-helix on one side, and a long loop.^[6-9]^

In response to frequent viral attacks, bacteria and archaea have evolved a wide array of anti-phage defense systems to safeguard themselves from phage infection,^[10, 11]^ including well-known mechanisms such as restriction-modification (RM), CRISPR-Cas, and abortive infection systems.^[12, 13]^ Among these, the abortive infection strategy is employed by systems like the defense-associated sirtuin (DSR) systems.^[14, 15]^ DSR2, a sirtuin (Sir2) domain-containing protein, demonstrates strong anti-phage defense capabilities through the abortive infection pathway.^[14]^ Previous research has shown that DSR2 can be activated by the newly synthesized TTP of the SPR phage, thus initiating the NAD+ depletion.^[14, 16-18]^ This activation occurs through the direct binding of TTP to DSR2, which triggers the enzymatic activity of the SIR2 domain, leading to the depletion of intracellular NAD+ and resulting in abortive infection.^[19]^ Upon injection of phage DNA into the bacteria and subsequent synthesis of phage proteins, the newly produced TTP monomer binds to the DSR2 tetramer, activating DSR2’s NADase activity. This reduction in cellular NAD+ levels leads to the death of the bacteria and prevents phage replication.^[20]^

SPR phages, which belong to the Siphoviridae family, are characterized by long, non-contractile tails and icosahedral heads.^[21, 22]^ These phages typically infect B. subtilis CU1050 and play roles in various biological processes, including horizontal gene transfer and bacterial population control.^[21]^ As noted, newly translated TTP of the SPR phage can interact with DSR2 to trigger an abortive infection response, allowing host cells to defend against phage SPR.^[14, 16-18]^ TTP also plays a crucial role in tube assembly.^[23, 24]^ Typically, TTPs form hexameric rings with a central tunnel, which are essential for the efficient delivery of the phage genome to host cells.^[25, 26]^ However, in some phages, such as T5, TTP can assemble into trimeric rings.^[23]^ In our study, we utilized cryo-EM to uncover a novel assembly mechanism of TTP from phage SPR. We discovered that TTPs in the SPR phage can not only assemble into a hexameric but also into a pentameric ring. Moreover, we revealed that at the end of the TTP tube, the β3 and β4 strands of each TTP shift toward the center of the tunnel, effectively blocking it. This structural rearrangement contrasts with the pentameric rings located at the center of the tube, where such blockage does not occur.

## Results

### Overall structure of the pentameric rings

To investigate the polymer structure of the phage tail tube proteins (TTP), we recombinantly expressed the TTP in the *E. coli* cells and subsequently performed the cryo-electron microscopy (cryo-EM) experiments on the purified sample. The cryo-EM images revealed that TTP self-assembles into tube-like polymers (Figs. 1). Notably, during the 2D classification step, we identified previously unseen TTP polymers composed of pentameric rings with a central tunnel. The ratio of these polymers to hexamers is approximately 1:10 based on the top-down views (Figs. 1). The 2D class averages indicated that the ends of these polymers had weak densities, suggesting structural flexibility. To improve the resolution of the TTP polymer structure, we designed a mask to focus on three layers of the polymers. After thorough 3D classification, we successfully resolved a cryo-EM map of three-layered TTP pentameric rings (TLTPR), in addition to the expected hexamers (Fig. 1, Figs. 1). The overall resolution of the TLTPR was 3.5 Å, with a local resolution of approximately 3 Å at the TLTPR’s center (Figs. 3(a)-(c), Table S1). The cryo-EM map of the TLTPR displayed clear side-chain densities, enabling us to accurately model the TLTPR structure (Figs. 4). Based on previous research (PDB: 8XKN),^[18]^ we divided the TTP monomer into two domains: domain 1 and domain 2, containing fifteen β-strands and one α-helix (Figs. 2(a)(b)).

The cryo-EM map of the TTP polymer reveals a tower-like structure with TTP pentamers stacked upon one another (Fig. 1(a)(b), 1(d)(e)). The TLTPR has a height of approximately 153 Å, with a width and length of about 124 Å (Fig. 1(a)(b), 1(d)(e)). The central tunnel has the diameter of around 39 Å, which is sufficient to accommodate the 2 nm double-stranded DNA (Fig. 1(c) and 1(f)). Notably, the angles between adjacent layers are different: the angle between the 1st and 2nd layer is 26°, while the angle between the 2nd and 3rd layer is 34°, resulting in a bent tube structure (Fig. 1(d)). We then plotted the electrostatic surface of the tube, and the result shows a highly negatively charged surface within the tunnel, suggesting it is favorable for DNA to pass through (Figs. 2(c)(d)).

**Fig. 1.**
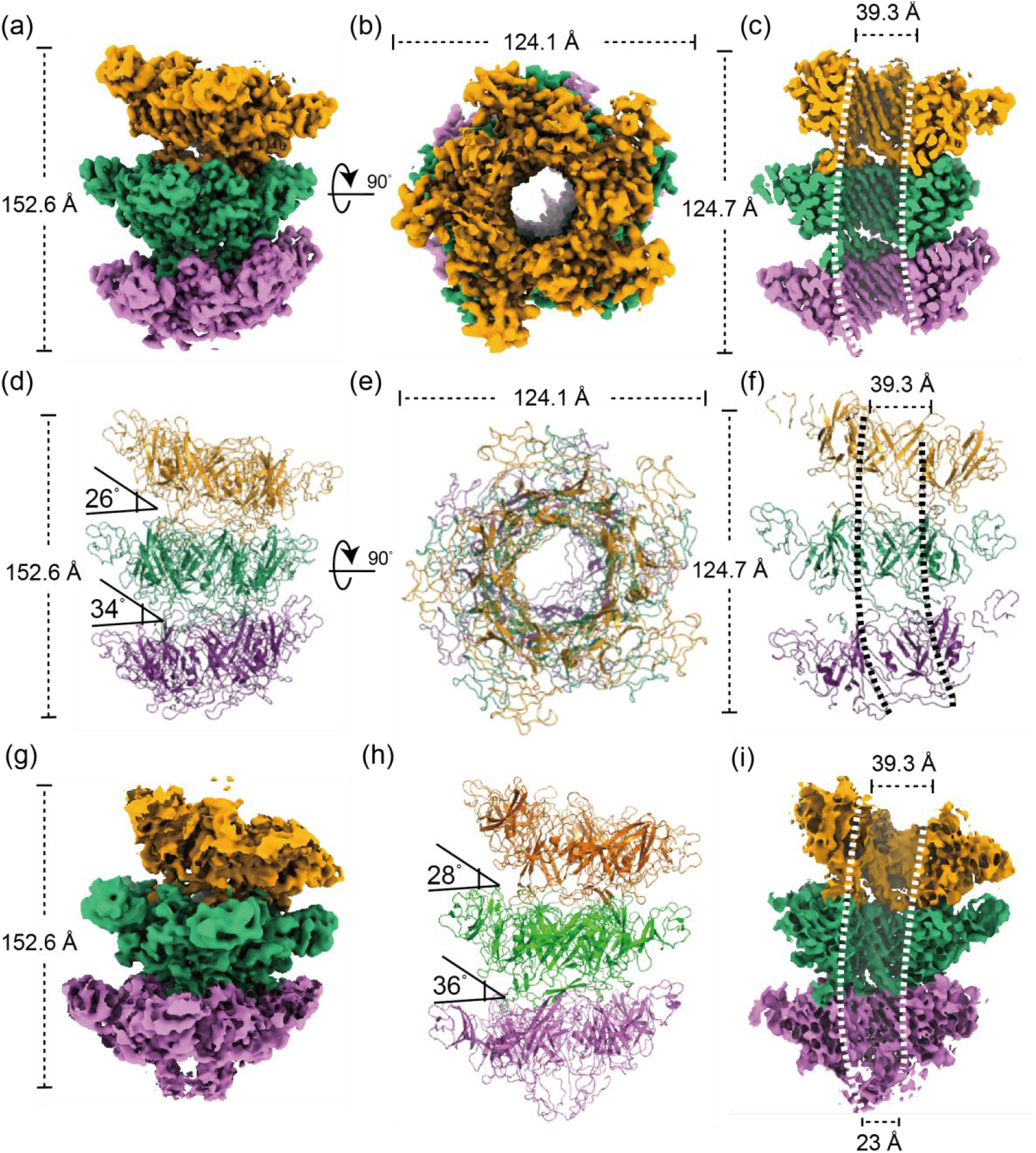
Overall Structure of the TLTRP. |(a), (b)| Cryo-EM maps of TLTRP shown in side (a) and top down (b) views. The corresponding structures are shown in |(d), (e)| The length, width and height are denoted in Å. |(c), (f)| Sagittal ‘slice’ view the cryo-EM map (c) and the corresponding structure (f), showing that the tunnel in the middle is bent at a certain angle. The dash line indicates the curvature of the tunnel. The diameter of the tunnel is denoted in Å. |(g)| Cryo-EM map of TLTRPE in side view(a). |h| The structure of TLTRPE. |(i)| Sagittal ‘slice’ view the cryo-EM map (g).

During our cryo-EM data analysis, we noted variability in the tube length. We hypothesized that the conformation of the pentamer at the TLTPR’s end (TLTPRE) might be affecting the elongation of the tube. To resolve the structure of TLTPRE, we first matched these TLTPR particles back to the micrographs and then manually adjusted the centers of the particles to align with the tube’s endpoint (Figs. 1; See Methods for details). By doing so, we successfully resolved the structure of TLTPRE at a resolution of 3.7 Å (Fig. 1(g)-(i), Figs. 2(e)(f), Figs. 3(d)-(f)). Further analysis demonstrated that the overall structure of the monomer in TLTPRE and TLTPR exhibits high similarity, with a calculated RMSD of 0.403 Å (Figs. 2(g)). However, we still discovered that the β3 and β4 strands in domain 1 of TLTPRE undergo significant conformational changes. Compared with TLTPR, the β3 and β4 strands in TLTPRE were shifted about 25 Å toward the center of the tube, thereby blocking the tunnel (Figs. 2(g)). Notably, we didn’t capture any other conformations at the tube’s end.

**Fig. 2.**
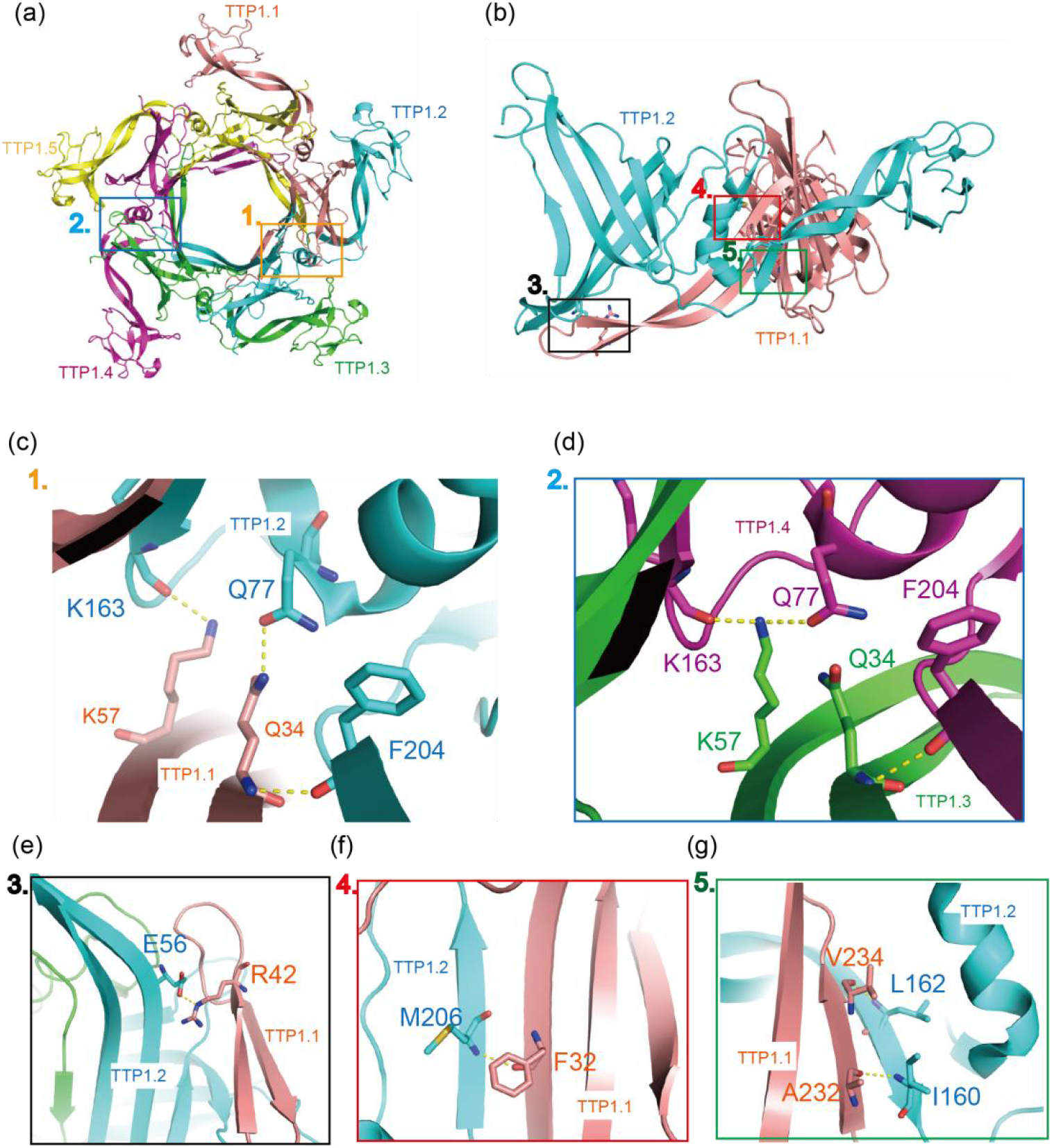
Interactions of Each TTP Protomers in TLTPR. (a) Isolation of the pentameric ring in TLTPR with numbers marked next to the subunit. The colored boxes are enlarged in (c) and (d). (b) Isolation of TTP1.1 and TTP1.2 The colored boxes are enlarged in panel (e), (f) and (g), respectively. Putative interactions are indicated by yellow dashed lines. Nitrogen, oxygen and sulfur are colored in blue, red and yellow, respectively.

### Interactions in one pentameric TTP ring

Comprehensive structural analysis revealed that all TTPs within each layer of the pentamer exhibit high structural similarity. To facilitate the structural analysis, we designated the subunits in the pentamer as shown in Fig. 2(a). We observed that neighboring monomers are stabilized by multiple hydrogen bonds, salt bridges, and extensive hydrophobic interactions (Fig. 2). The interface areas between these two adjacent monomers are measured approximately 13,037 Å^2^. We discovered that the N-terminus of TTP1.1 interacts with both the N- and C-termini of the adjacent TTP1.2, which contributes to the stabilization of the pentamer (Fig. 2(b)). The interactions between monomers within the pentameric ring showed a high degree of similarity, with several common interactions identified (Fig. 2(b)). For instance, hydrogen bonds were observed between K163, F204, E56, M206, I160, and L162 on TTP1.1, and K57, Q34, R42, F32, A232, and V234 on TTP1.2 (Fig. 2(c), 2(e)-(g)). However, we also noted subtle differences between the monomers in the pentamer ring. For example, while hydrogen bonds were formed between Q34 on TTP1.1 and Q77 on TTP1.2 (Fig. 2(c)), the corresponding interactions differ between TTP1.3 and TTP1.4 (Fig. 2(d)). Specifically, the hydrogen bond between Q34 on TTP1.3 and Q77 on TTP1.4 was disrupted, and instead, K57 on TTP1.3 forms two interactions with Q77 and K163 on TTP1.4 (Fig. 2(d)). We proposed that these interactions between monomers probably contribute significantly to the stabilization of the TLTPR.

### Comparison of the pentameric and hexameric TTP rings

Due to the absence of one TTP subunit, the diameter of the central tunnel decreased from approximately 50 Å in the hexameric ring to 40 Å in the pentameric ring (Fig. 3(a)). Moreover, the measured distances between the centers of mass (COM) of the subunits indicate a pseudo-symmetry within the pentamer (Fig. 3(b)). Unlike the hexamer, which exhibits C6 symmetry, the distances between the COMs in the pentamer display variability, ranging from 27 to 28 Å, compared to the more uniform distance of approximately 29.7 Å observed in the hexamer (Fig. 3(b)).

**Fig. 3.**
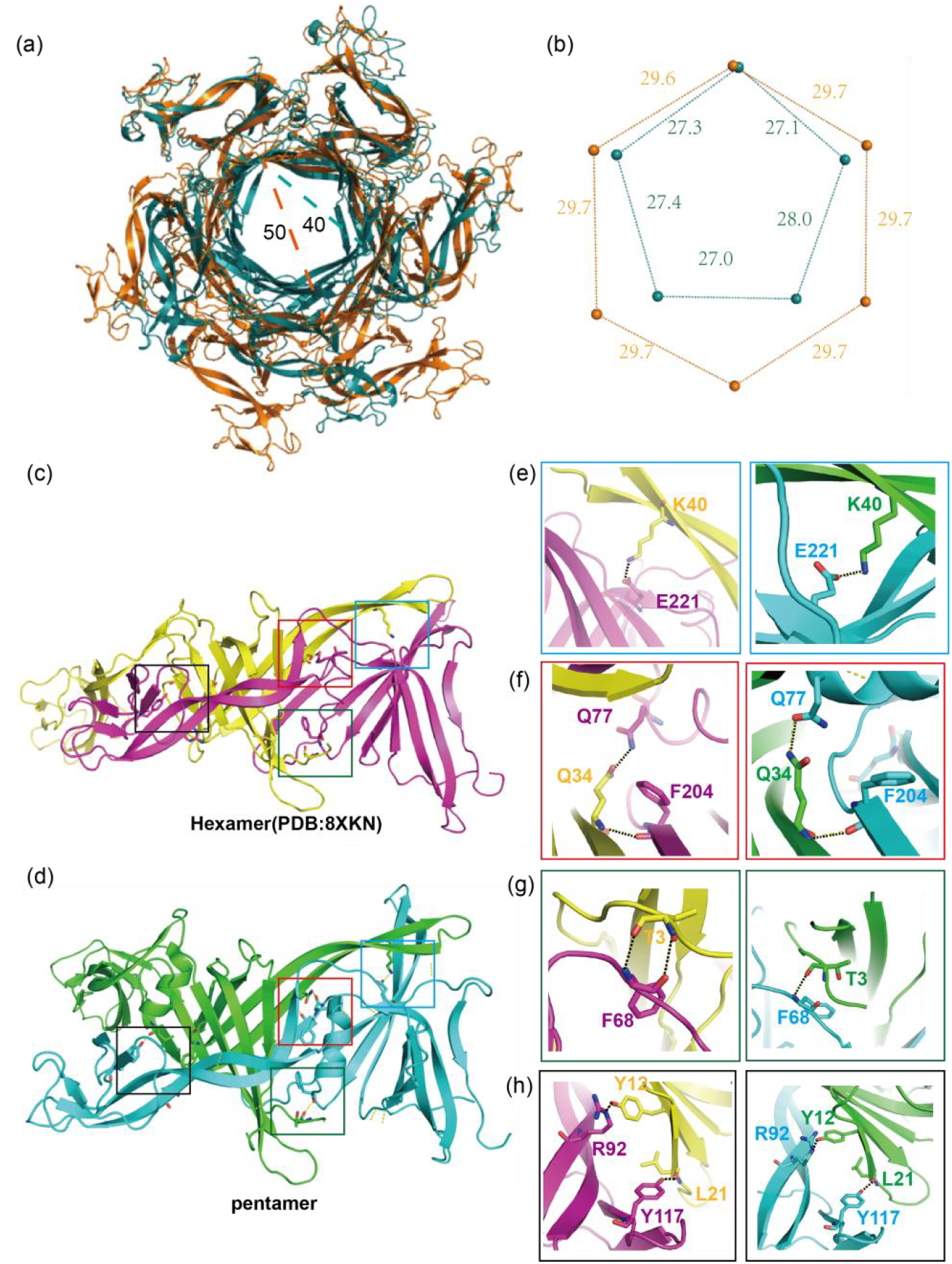
Same Interactions between Pentameric and Hexameric rings. (a) Alignment of the pentameric and hexameric (PDB: 8XKN) rings. The pentameric and hexameric ring are in orange and green, respectively. (b) Schematic illustrating the distances (Å) between the centers of mass (COM) of the subunits in the pentameric and hexameric rings. The color code is the same to panel (a). |(c), (d)| Isolation of the two adjacent subunits from pentameric (c) and hexameric rings (d). The boxed areas are enlarged in panel (e) to (g) with the same color code. Putative interactions are indicated by black dashed lines. Nitrogen and oxygen are colored in blue and red, respectively.

To further investigate the interactions within the TTP assemblies, we isolated two adjacent subunits from both the hexameric (PDB: 8XKN)^[18]^ and pentameric rings for detailed structural analysis (Fig. 3(c)(d)). Our findings revealed that the N-terminus of the TTP forms extensive interactions with the neighboring subunit in both ring types (Fig. 3(e)-(h)). Interestingly, many of these interactions remained consistent during the transition from hexamer to pentamer, highlighting their potential importance in maintaining structural integrity. Specifically, we identified several key hydrogen bonds between the adjacent subunits, including interactions between Q34 and Q77, Q34 and F204, F68 and T3, R92 and Y12, and Y117 and L21 (Fig. 3(e)-(h)). Additionally, we observed a salt bridge between E221 and K40, further stabilizing the interface between the subunits (Fig. 3(e)). These conserved interactions suggest their critical roles in the structural assembly of both hexameric and pentameric TTP rings.

The interactions unique to the pentameric structure likely play crucial roles in the formation of pentameric rings (Fig. 4). The pentameric TTP adopts a more compact conformation compared to its hexameric counterpart. Our analysis suggests that the formation of pentameric rings involves both the disruption of certain interactions present in the hexameric rings and the establishment of new ones (Fig. 4(a)(b)). For example, hydrogen bonds observed in the hexameric rings, such as those between K125 and D18, F90 and E174, Q34 and G202, and M231 and I162, are disrupted in the pentameric assembly (Fig. 4(e)-(h)). Conversely, the pentameric rings exhibit new hydrogen bonds, including those between E236 and R158, R92 and D10, K88 and Q194, K57 and K163, and K57 and Q77 (Fig. 4(c)-(j)). These alterations predominantly occur in the β3, β4, and β10 strands, highlighting the importance of these β strands in the formation and stabilization of the pentameric rings (Figs. 2(a)).

**Fig. 4.**
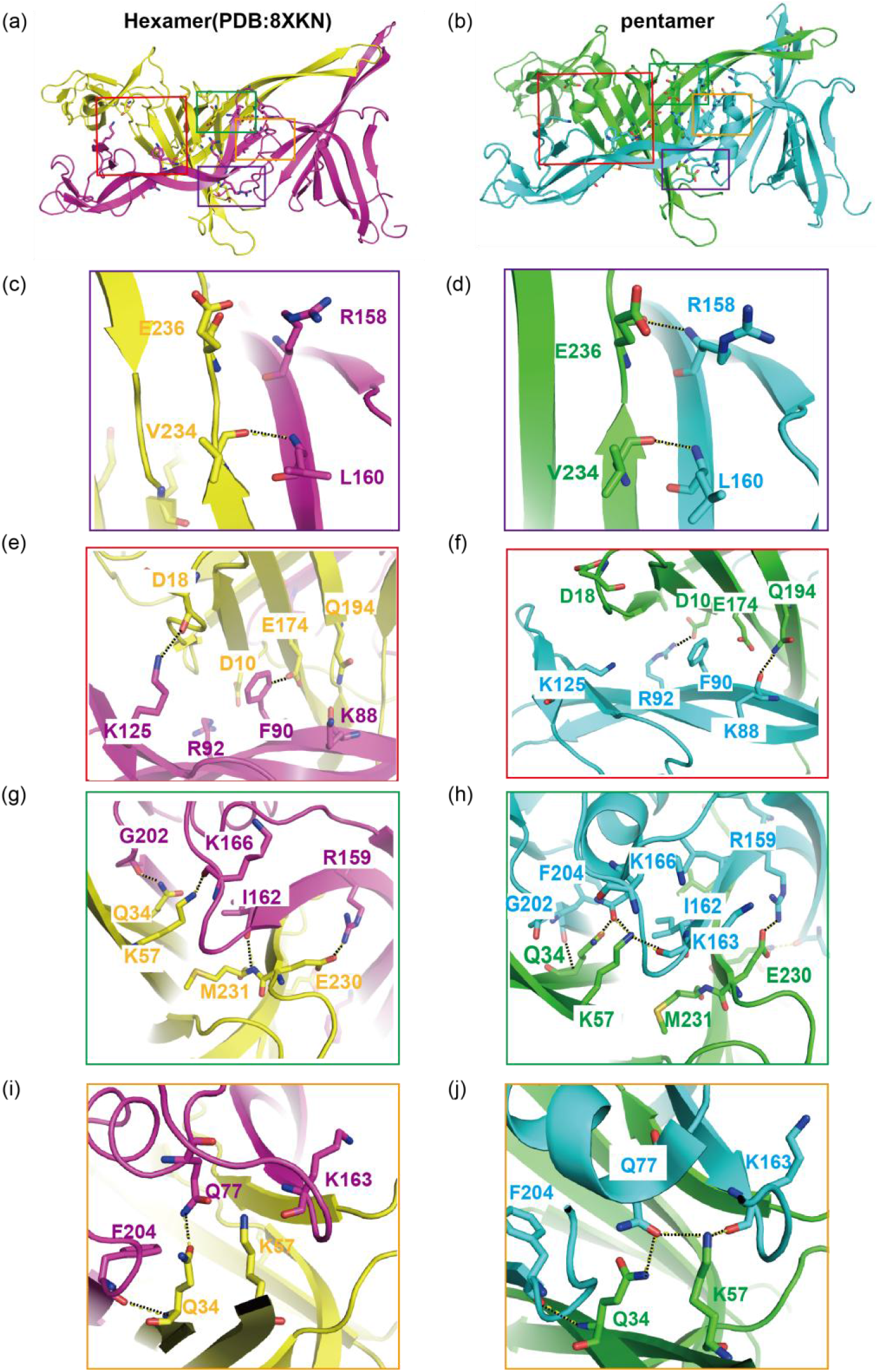
Comparison of Interactions Between the Pentameric and Hexameric ring. |(a), (b)| Isolation of two adjacent subunits from the pentameric (a) and hexameric (b) ring, respectively. The interfaces showing different interactions are marked by boxes in different colors. The corresponding areas are enlarged in panel (c) to (g) following the same color code. Putative interactions are indicated by black dashed lines. Nitrogen and oxygen are colored in blue and red, respectively.

### Interactions between the stacked pentameric layers

To elucidate the interactions between the pentameric layers, we assigned numerical identifiers to each TTP subunit within the polymer (Fig. 5(a)). The buried interface area between the first and second layer measures 28,813 Å^2^, which is greater than the interface area between the second and third layers, measuring 28,189 Å^2^ (Fig. 5(b)). Our observations revealed distinct interactions between the pentameric layers that contribute to the curvature of the tube structure. Our structural analysis indicated that fewer interactions were present on the looser left side of the structure (Fig. 5(b), Figs. 5(a)). Specifically, residues R42, G44, and N47 on TTP1.1 form multiple hydrogen bonds with residues on TTP2.2 (Q28 and A27) and on TTP2.3 (N210 and Q211) (Fig. 5(c)). In contrast, these interactions are entirely absent regarding to TTP2.1 (Fig. 5(d)). Additionally, an unexpected hydrogen bond was observed between I52 on TTP2.1 and E185 on TTP3.1 (Fig. 5(f)), whereas the corresponding interaction between TTP1.1 and TTP2.1 was not present (Fig. 5(e)).

**Fig. 5.**
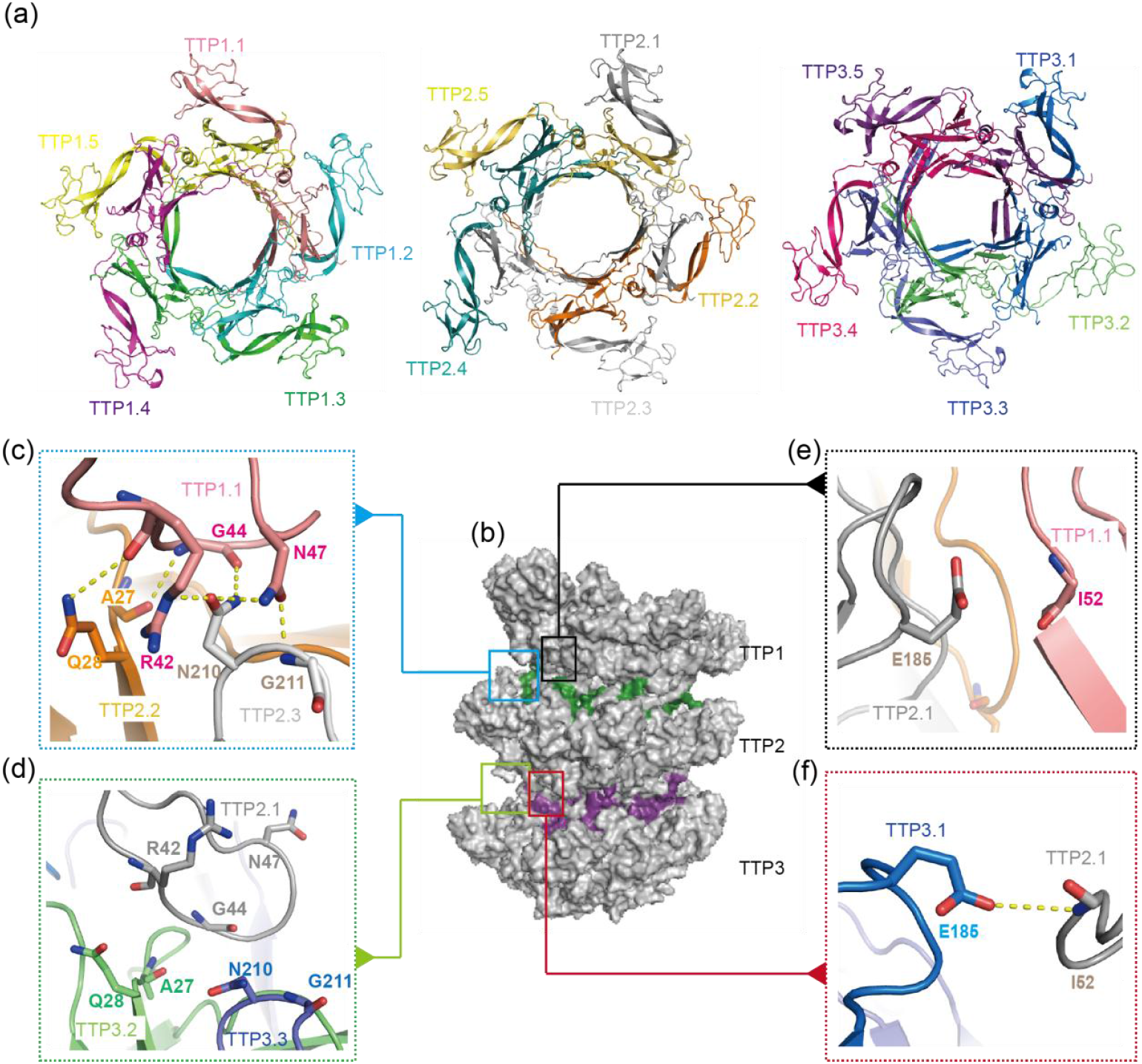
The Interaction Interfaces Between Stacked layers in the TLTPR. (a) Isolated layers are shown with the corresponding number labeled next to each subunit. (b) TLTPR is shown in surface representation. The green and purple areas indicate the interfaces between the first and second layer, and between second and third layer, respectively. The boxed areas are enlarged in panel (c) to (f) following the same color code. Putative interactions are indicated by yellow dashed lines. Nitrogen and oxygen are colored in blue and red, respectively.

In comparison to the left side, the right side of the structure exhibits a greater number of interactions (Figs. 5(a)), particularly in the loop connecting β3 and β4 on TTP1.4 and TTP2.4 (Figs. 2(a), Figs. 5(b)-(d), 5(h)). Significant interactions were observed between adjacent layers, involving residues Q6, T8, A27, G46, Y51, K54, K146, R170, E185, N210, and D227 (Figs. 5(b)-(k)). Notably, between the first and second layers, R170 on TTP1.4 forms interactions with P182 and E185 on TTP2.3 (Figs. 5(g)). However, these interactions are reduced between TTP2.4 and TTP3.3 (Figs. 5(j)). Additionally, hydrogen bonds involving TTP1.4 were found between residue G44 of TTP1.4 and residues T8 and A27 of TTP2.5 (Figs. 5(d)), yet the corresponding interactions are lost at the interface between TTP2.4 and TTP3.5 due to conformational changes in the loop between β3 and β4 (Figs. 5(h)). At the interface between the TTP1.4 and TTP2.5, the D227 interacts only with K146(Figs. 5(f)), while at the interface between TTP2.4 and TTP3.5, hydrogen bonds are formed between D227 and both K146 and H94 (Figs. 5(i)). Interestingly, hydrogen bonds are formed between TTP3.5 (A27 and Q28) and TTP2.4 (G46 and N47), while A27 and Q28 on TTP2.5 form hydrogen bonds with G44 and I45 on TTP1.4 (Figs. 5(h) and Figs. 5(d)).

## Discussion

In this study, we investigated the assembly mechanism of TTP and identified two novel assembly states TLTPR and TLTPRE. In addition to the conventional hexameric rings, our cryo-EM maps reveal that TTP can assemble into pentameric rings, resulting in a tube with a 4 nm central tunnel (Fig. 1(c)). This finding suggests a specialized assembly mechanism distinct from SPR phages. Compared to the TTP hexameric structure (PDB: 8XKN),^[18]^ we identified specific interactions within the pentameric rings which are probably crucial for their assembly. Moreover, analysis of interactions between different layers of the tube showed significant variations that contribute to the curvature of the tube. The SPR phage has a long, flexible, non-contractile tail; due to the phage’s long tail, a perfectly straight and rigid tail would require more space and could limit its ability to invade host cells effectively. Having a curved or flexible tail might make the phage more adaptable, helping it to better protect its genome and more easily infect host cells. This curvature may impact the trajectory of DNA during ejection, potentially adapting the tube structure for optimized phage-host interactions. At the end of the tail tube, we observed that two β-strands from TTP subunits block the central tunnel (Figs. 2(g)), potentially regulating DNA ejection by preventing premature release and ensuring controlled infection.

Recent insights into bacterial anti-phage defense mechanisms have highlighted the importance of specific amino acids in the TTP of the SPR phage. TTP plays a critical role during phage infection, particularly through its interaction with Defense-associated Sirtuin 2 (DSR2), which is vital to the bacterial defense strategy. Notably, previous research demonstrated that the loop region of TTP (amino acids 202-217), especially residues F204 and M206, inserts into a hydrophobic pocket on DSR2 to stabilize the interaction^[16]^. Interestingly, our structural analysis revealed that F204 and M206 participate in interactions between adjacent subunits, underscoring their significance (Fig. 2(c)-(d) and Fig. 2(f)).

Previous research indicated that mutations in amino acids located in the loop between β3 and β4 can inhibit the formation of the DSR2-TTP complex, thereby compromising bacterial defense against phage infection.^[16]^ This loop, encompassing amino acids 35-55, has been shown to contribute to strong hydrophobic interactions with DSR2.^[16]^ Interestingly, our study found that this loop is crucial for connecting the pentamers (Fig. 5(c) and Fig. 5(f)). Specifically, our investigation of TLTPRE revealed that the central tunnel is obstructed by the loop between β3 and β4 (Figs. 2(g)). We speculated that the exposed β3 and β4 might help TTP bind to other partners like DSR2; however, we cannot exclude the possibility that they may recruit more TTP pentameric rings to facilitate the elongation of TLTPR.

We discovered a novel TTP assembly state and provided us a critical new insight into the structural biology of bacteriophages, highlighting the diversity of TTP assembly mechanisms. However, we cannot exclude the possibilities that the pentameric assembly might result from the variations in protein expression, purification, and cryo-EM sample preparation.^[27-29]^ While we have resolved the cryo-EM structures of a novel TTP assembly state, it remains unclear whether the formation of pentameric rings is a component of an anti-phage defense mechanism or a naturally occurring process in phage tail assembly. Further studies are necessary to elucidate the role of pentamer formation within the broader context of tail tube assembly and to explore its potential implications for phage defense strategies.

## Methods

### Protein expression and purification

The tail tube protein (TTP; WP:010328117) gene from *Bacillus subtilis* bacteriophage SPR was synthesized by GenScript and subsequently cloned into the pETDuet-1 vector. To facilitate protein purification, a C-terminal 6×His tag was fused to the TTP gene. Protein expression was carried out in *Escherichia coli* BL21 (DE3) cells. When the optical density at 600 nm (OD600) of the cells in LB medium reached approximately 0.8, protein expression was induced by adding 0.25 mM isopropyl β-D-1-thiogalactopyranoside (IPTG) and allowed to proceed overnight.

For protein purification, the overnight cultures of BL21 (DE3) cells were harvested by centrifugation, and the resulting cell pellets were resuspended in lysis buffer (40 mM Tris-HCl, pH 8.0, 10 mM imidazole, 150 mM NaCl, and 1 mM phenylmethylsulfonyl fluoride [PMSF]). The cells were lysed using an ultrasonic cell disruptor, followed by centrifugation at 25,000g for 50 minutes at 4°C to remove cell debris. The supernatant was subsequently applied to a 5 mL Ni-NTA affinity column (QIAGEN) via gravity flow. After washing with lysis buffer, the His-tagged TTP protein was eluted with a buffer containing 300 mM imidazole. Further purification of the TTP protein was accomplished using a HiTrap Q HP ion-exchange column (GE Healthcare), with gradient elution with NaCl buffer. The eluate fractions containing the target protein were then concentrated and loaded onto a size exclusion column, Superdex 200 10/300 (Cytiva), which was pre-equilibrated with buffer containing 20 mM Tris-HCl, pH 8.0, and 100 mM NaCl.

### Cryo-EM sample preparation and data collection

The 300 mesh R1.2/1.3 Quantifoil carbon grids (Au, Electron Microscope Sciences) were glow discharged using a PELCO easyGlow (0.39 mBar air) glow discharge instrument at 15 mA for 60 seconds. Using a Vitrobot Mark IV (FEI), 3 μl of HEPES buffer containing 3mg/mL TTP protein was applied to the grids, which were subsequently blotted with filter papers at a blot force of 4 for 7 seconds in 100% humidity at 4 °C. The sample was then rapidly frozen by plunging into liquid ethane, pre-cooled by liquid nitrogen. Cryo-EM movies were recorded using a 300kV Titan Krios electron microscope (Thermo Fisher) equipped with a K3 Summit direct electron detector. A slit width of 20 eV was used, and images were captured at a super-resolution pixel size of 0.415 Å. Movies were automatically collected by SerialEM. The total electron dose used was 66 electrons/Å^2^, evenly distributed across 32 frames. Defocus values ranged from -1.5 to -2.5 μm.

### Cryo-EM data processing

All movies were initially aligned and dose-weighted using MotionCor2,^[30]^ and defocus values were estimated with CTFFIND4.^[31]^ Particle picking was performed automatically using CryoSparc.^[32]^ A total of 2,407,982 raw particles were selected for subsequent 2D classification. After 2D classification, 555,749 particles were retained for a round of Ab-Initio reconstruction with four classes. One of these classes, containing 149,568 particles, was selected for further heterogeneous refinement. And 49,339 particles were retained for one round of non-uniform refinement, resulting in a map with a resolution of 3.5 Å. To resolve the structure of the TTP pentamer at the ends of the filament, approximately 200 particles from the 49,339 good particles were re-extracted and matched back to the micrographs. The centers of these particles were then manually adjusted, and particle picking was subsequently performed using Topaz,^[33]^ yielding 362,574 particles. After one round of 2D classification, particles exhibiting a blocked end were selected for further 3D classification. Finally, 96,299 particles were used for a round of non-uniform refinement, producing a map with a resolution of 3.7 Å. The TLTRP and TLTRPE were sharpened using DeepEMhancer^[34]^ and Locscale,^[35]^ respectively. Figures were prepared with PyMOL and UCSF ChimeraX.^[36]^ The centers of mass (COM) of each TTP protomer in the pentamer and hexamer and the electrostatic surface of the pentamer are calculated by pymol.

### Model building

For the models of the pentamers structure of TTP, we first generated a homologous model of TTP using AlphaFold2.^[37]^ To build the model of TTP polymer, we first built the model of the TTP monomer. Due to the flexibility of the TTP monomer, we separated the homologous model into domain 1 and domain 2. Then we manually fit domain 1 and domain 2 to the TTP monomer in Chimera,^[38]^ followed by manual refinement in Coot.^[39, 40]^ Phenix^[41, 42]^ real space refinement was then performed to further refine the TTP monomer against the TTP map. After obtaining the model of the TTP monomer, we next fit 15 copies of the TTP monomers into the TTP polymer map in Chimera, followed by a round of phenix real space refinement. The final CC value was 0.74. Similarly, to build the model for the end-blocked TTP polymer, we first modeled the TTP monomer at the end of the filament. We then replaced the five monomers blocking the tunnel in Chimera. Finally, a round of phenix real space refinement was performed to refine the model against the cryo-EM map. The final CC value was 0.75. The interactions in the structure were analyzed using PDBePISA.^[43]^

## Data and code availability

The atomic models of a pentameric assembly of the tail tube protein (TTP) and a pentameric assembly of the tail tube protein (TTP) at the tube’s end have been deposited into the PDB with accession codes 9JGI and 9JGH, respectively. The cryo-EM density maps have been deposited into the Electron Microscopy Data Bank under the accession codes EMD-61465 and EMD-61464, respectively.

### Acknowledgments

We thank D. Sun and Y. Wang at the Cryo-EM Facility at the Institute of Physics, Chinese Academy of Sciences (IOP, CAS), and X. Huang, B. Zhu, X. Li, L. Chen, and other staff members at the Center for Biological Imaging (CBI), Core Facilities for Protein Science at the Institute of Biophysics, Chinese Academy of Science (IBP, CAS), for their support in cryo-EM data collection. This work was supported by grants from the Chinese Academy of Sciences (CAS) (E4V4061 and E2VK311).

## Author Contributions

K.S., X.G., S.C. and H.Z. designed the project. L. W., Y. H., K. S. and K. Z. performed cryo-EM sample preparation, cryo-EM data collection, and the cryo-EM data analysis. L.W., K. S. and Y. H. performed the model building and structure analysis. L. W., Y. H., K. Z., X. G., K. S., S. C. and H. Z. wrote the manuscript.

