## Supplemental Figure and Table for "Architecture of a pentameric assembly of the tail tube protein in SPR phages"

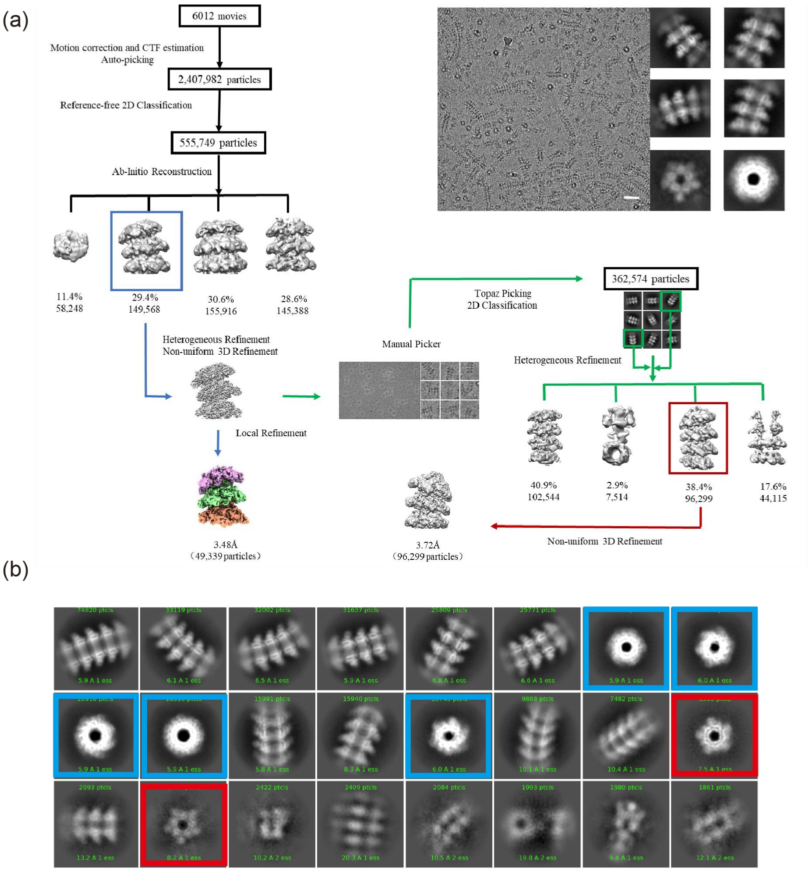


**Figs. 1. Cryo-EM data processing workflow of tail tube protein (TTP).**

(a) The workflow for Cryo-EM data processing of the tail tube protein (TTP).

(b) Typical 2D classifications of TTP, where the blue boxes highlight the standard top-down view of hexamers, and the red boxes highlight the standard top-down view of pentamers.

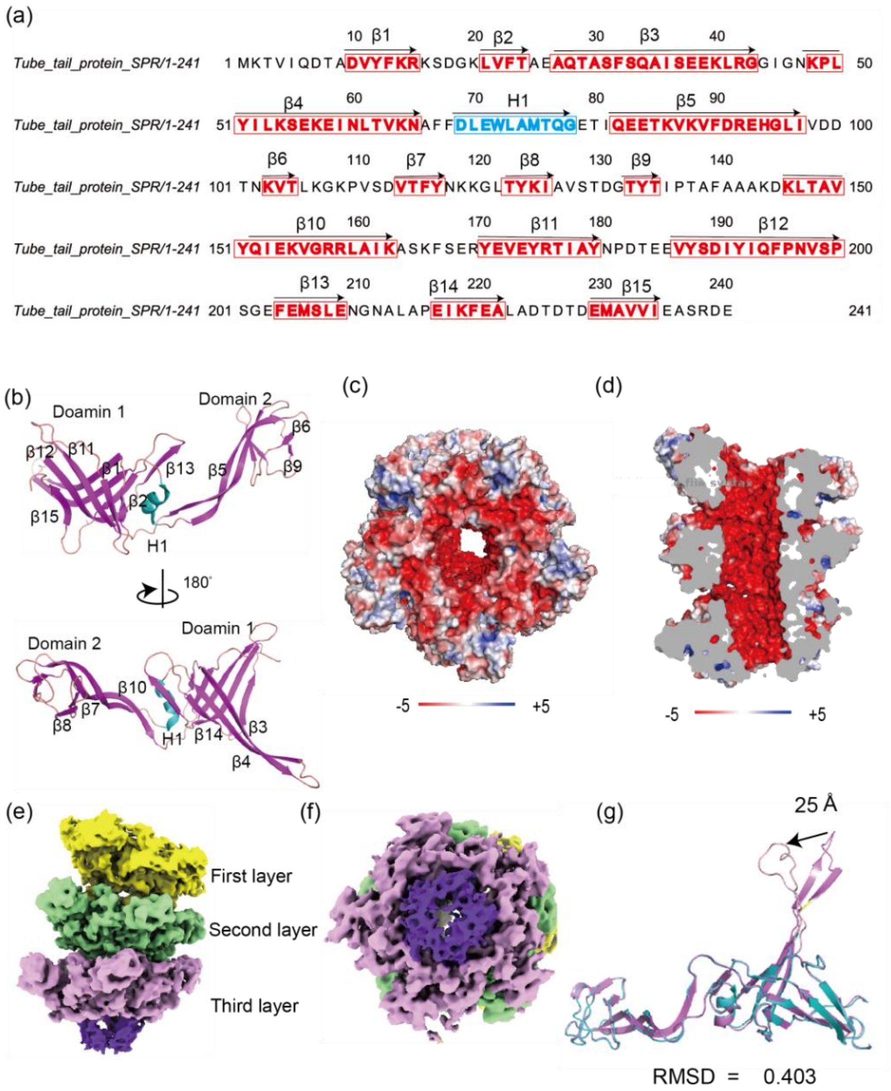


**Figs. 2. The secondary structure and electrostatic surface of the pentameric rings.**

(a) The sequence of the tail tube protein, with key features marked on the sequence. The residues in red correspond to the β-sheets, while the residues in blue represent the α-helix. The numbering of the β-sheets and α-helix is marked. (b) Overall structure of the TTP monomer. The TTP monomer is divided into Domain 1 and Domain 2 with the secondary elements labeled. (c) Electrostatic surface of the pentameric rings. (d) Sagittal 'slice' view along the pore axis of the electrostatic surface of the pentameric rings. |(e), (f)| Cryo-EM structure of the TTP polymer at the tube's end shown in side view (e) and topdown view (f). The structure consists of three layers, each in a different color. (g) Alignment of the TTP monomers in the 2nd and 3rd layer of the pentamer. The monomer in 2nd and 3rd layer are colored in blue and purple, respectively. The loop between β3 and β4 is labeled light pink in the 3rd layer, and there is a 25 Å shift compared with the loop between β3 and β4 in the 2nd layer, labeled in purple.

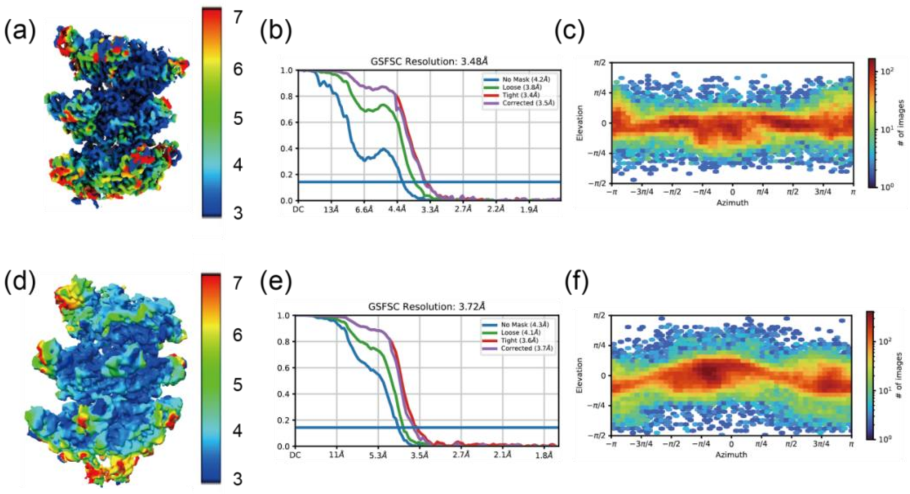


**Figs. 3. Cryo-EM Structure Validation of the Pentamer.**

(a) Local resolution map for TLTPR, (b) FSC curves for the cryo-EM reconstruction of TLTPR, and (c) particle orientation distribution for TLRPR.

(d) Local resolution map for TLTPRE, (e) FSC curves for the cryo-EM reconstruction of TLTPRE, and (f) particle orientation distribution for TLTPRE.

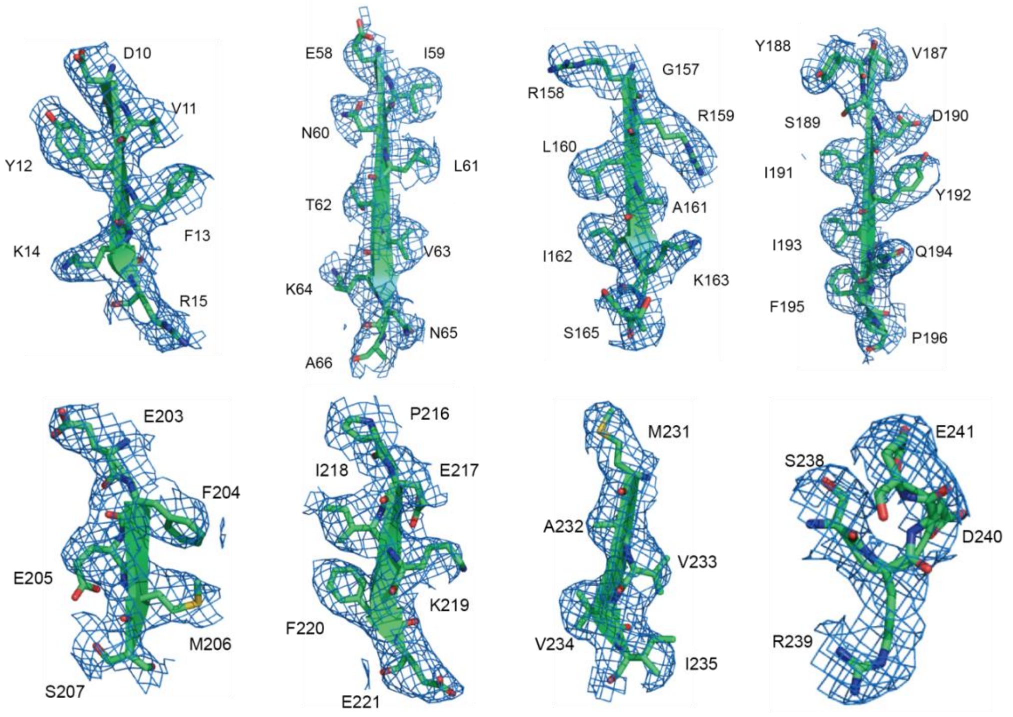


**Figs. 4. The representative density maps of the TTP protomer.**
The corresponding residues are labeled next to the density map.

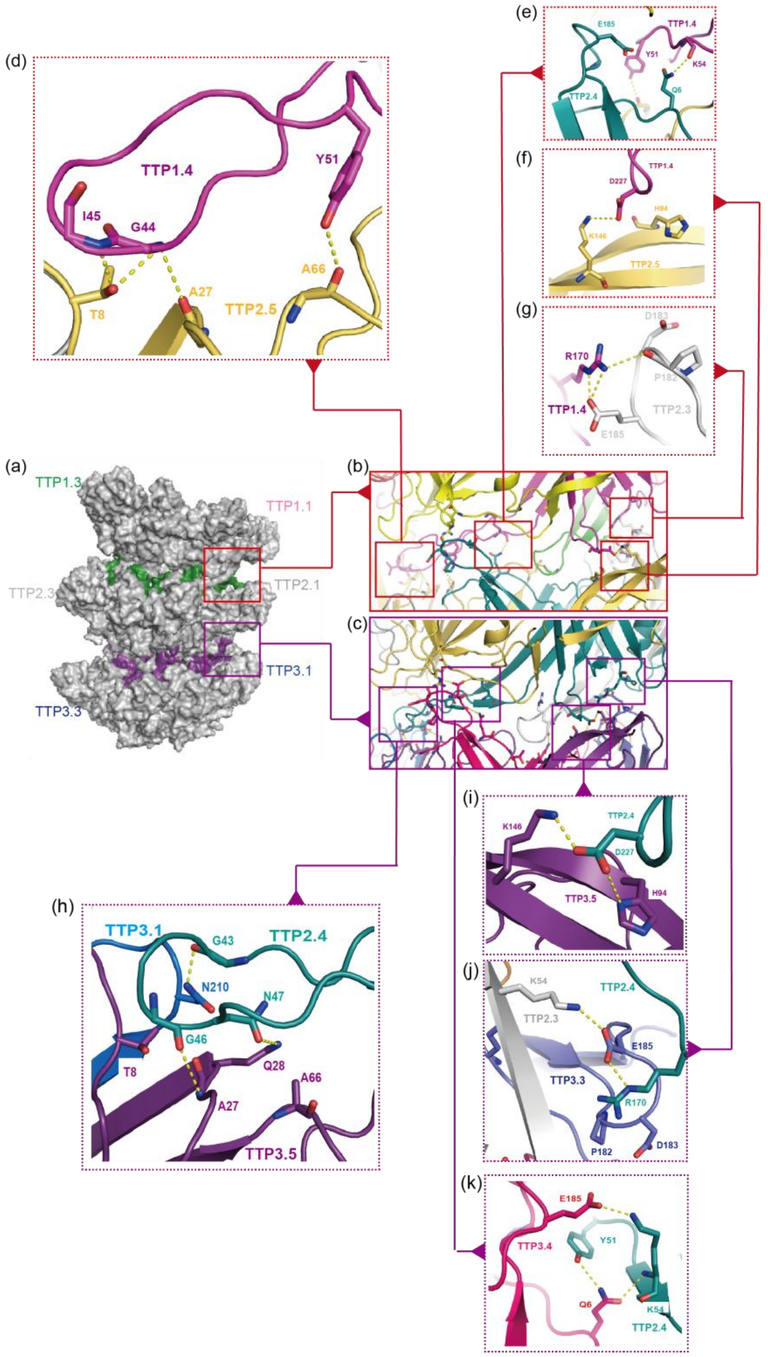


**Figs. 5. The Different Interaction Interfaces Between Layers in the Structure.** The interactions between each layer are marked by boxes in different colors (a). The interactions between the first and second layers are circled with red boxes. The interaction interfaces are enlarged in (b), where four interacting interfaces are separately marked by red boxes. These enlarged boxes with red dashed borders |(d)-(g)| show the interactions between specific amino acids corresponding to (b). The interactions between the second and third layers are indicated by purple boxes. The interaction interfaces are enlarged in (c), marked by four purple boxes. These enlarged boxes with purple dashed borders |(h)-(k)| show the interactions between specific amino acids corresponding to (c). The color code is the same to Fig. 5. Putative interactions are indicated by yellow dashed lines.

**Table S1 Cryo-EM data collection, refinement, and validation statistics**

| | TLTPR (EMD-61465, PDB 9JGI) | TLTPRE (EMD-61464, PDB 9JGH) |
|---|---|---|
| Magnification | 81000x | 81000x |
| Voltage (kV) | 300 | 300 |
| Electron exposure ($e^{-}$/Å$^2$) | 60 | 60 |
| Defocus range (μm) | -1.5 to -2.5 | -1.5 to -2.5 |
| Pixel size (Å) | 0.83 | 0.83 |
| Symmetry imposed | C1 | C1 |
| Initial particle images (no.) | 555,749 | 362,574 |
| Final particle images (no.) | 49,339 | 96,299 |
| Map resolution (Å) | 3.5 | 3.7 |
| FSC threshold | 0.143 | 0.143 |
| Map resolution range (Å) | 3-7 | 3-7 |
| **Refinement** | | |
| Initial model used (PDB code) | AlphaFold | 9JGI |
| Model resolution (Å) | 3.8 | 4.4 |
| FSC threshold | 0.5 | 0.5 |
| Map sharpening Method | DeepEMhancer | Locscale |
| **Model composition** | | |
| non-hydrogen atoms | 27,798 | 28,590 |
| Protein residues | 3,513 | 3,615 |
| Protein | 27,798 | 28,950 |
| **R.m.s. deviations** | | |
| Bond lengths (Å) | 0.005 | 0.021 |
| Bond angles (°) | 1.119 | 1.267 |
| **Validation** | | |
| MolProbity score | 1.82 | 2.07 |
| Clashscore | 5.07 | 7.76 |
| Poor rotamers (%) | 0.89 | 0.93 |
| **Ramachandran plot** | | |
| Favored (%) | 89.89 | 85.94 |
| Allowed (%) | 9.30 | 13.33 |
| Disallowed (%) | 0.81 | 0.73 |
